# Reprogramming acetogenic bacteria with CRISPR-targeted base editing *via* deamination

**DOI:** 10.1101/2020.04.20.047845

**Authors:** Peng-Fei Xia, Isabella Casini, Sarah Schulz, Christian-Marco Klask, Largus T. Angenent, Bastian Molitor

## Abstract

Acetogenic bacteria are rising in popularity as chassis microbes in biotechnology due to their capability of converting inorganic one-carbon (C1) gases to organic chemicals. To fully uncover the potential of acetogenic bacteria, synthetic-biology tools are imperative to either engineer designed functions or to interrogate the physiology. Here, we report a genome-editing tool at a one-nucleotide resolution, namely base editing, for acetogenic bacteria based on CRISPR-targeted deamination. This tool combines nuclease deactivated Cas9 with activation-induced cytidine deaminase to enable cytosine-to-thymine substitution without DNA cleavage, homology-directed repair, and donor DNA, which are generally the bottlenecks for applying conventional CRISPR-Cas systems in bacteria. We designed and validated a modularized base-editing tool in the model acetogenic bacterium *Clostridium ljungdahlii*. The editing principles were investigated, and an *in-silico* analysis revealed the capability of base editing across the genome. Moreover, genes related to acetate and ethanol production were disrupted individually by installing premature STOP codons to reprogram carbon flux towards improved acetate production. This resulted in engineered *C. ljungdahlii* strains with the desired phenotypes and stable genotypes. Our base-editing tool promotes the application and research in acetogenic bacteria and provides a blueprint to upgrade CRISPR-Cas-based genome editing in bacteria in general.

**Significance:** Acetogenic bacteria metabolize one-carbon (C1) gases, such as industrial waste gases, to produce fuels and commodity chemicals. However, the lack of efficient gene-manipulation approaches hampers faster progress in the application of acetogenic bacteria in biotechnology. We developed a CRISPR-targeted base-editing tool at a one-nucleotide resolution for acetogenic bacteria. Our tool illustrates great potential in engineering other A-T-rich bacteria and links designed single-nucleotide variations with biotechnology. It provides unique advantages for engineering industrially relevant bacteria without creating genetically modified organisms (GMOs) under the legislation of many countries. This base-editing tool provides an example for adapting CRISPR-Cas systems in bacteria, especially those that are highly sensitive to heterologously expressed Cas proteins and have limited ability of receiving foreign DNA.

## Introduction

Global climate change is challenging the future of human societies, resulting in the need for a sustainable food supply and greener synthesis of fuels and chemicals. One possible solution is by applying biotechnology to convert inorganic one-carbon (C1) gases, such as carbon dioxide (CO_2_) and carbon monoxide (CO), into protein, biofuels, and commodity chemicals (1–4). Both gases are already available in large quantities, including in synthesis gas (syngas) and industrial waste gases (5). Many studies have found that the model acetogenic bacterium *Clostridium ljungdahlii* can convert these gases with hydrogen gas (H_2_) into mainly acetate and ethanol *via* the Wood-Ljungdahl pathway (1, 6). LanzaTech, Inc. (Skokie, IL, USA) has already completed the industrial scale-up by utilizing the closely related acetogenic bacterium (*Clostridium autoethanogenum*). However, synthetic biology and metabolic engineering are imperative to improve the productivity further and to expand the product spectrum (1, 3, 7, 8). Currently, the lack of efficient genome-editing tools delays the progress at the molecular level to optimize acetogenic bacteria for biotechnology.

Clustered regularly interspaced short palindromic repeats (CRISPR)-Cas-based genome editing is a RNA programmable, precise, and robust approach for gene perturbation, and has been applied in a plethora of living organisms (9), revolutionizing science. Recently, CRISPR-Cas systems were also adapted to be functional in acetogenic bacteria for gene deletion, insertion, and regulation. To date, different CRISPR-Cas systems (*e.g.*, Cas9 and Cas12a) have been established in *C. ljungdahlii* (10–12), *Eubacterium limosum* (13), and *C. autoethanogenum* (14). For these systems, first, the Cas protein (*e.g.*, Cas9 from *Streptococcus pyogenes*) is targeted to a highly specific site on the genome by a guide RNA (gRNA). Then, the Cas protein cleaves the genomic DNA at this site and introduces a double-strand break. To survive, the cell has to repair the double-strand break through DNA repairing mechanisms such as homology-directed repair or non-homologous end joining (15). For homology-directed repair, a donor DNA has to be provided as a template, which has to contain homologies to the genome on both sides of the double-strand break. Depending on the design of the donor DNA, it is possible to generate a variety of desired mutations such as point mutations, gene deletions, and gene insertions. The homology-directed repair of the double-strand break results in genome editing at the target site without leaving a selective marker (*i.e.*, antibiotic resistance gene) and scar.

However, CRISPR-Cas-based genome editing is generally challenging in bacteria, because the Cas nuclease is often toxic to bacteria, and bacteria typically lack efficient homology-directed repair or non-homologous end joining machineries to repair the double-strand break (16). Therefore, it is essential that a sufficient number of cells receive the CRISPR-Cas system to ensure that enough cells survive a DNA cleavage by undergoing the inefficient homology-directed repair process with donor DNAs (15). This renders the CRISPR-Cas system even more difficult for acetogenic bacteria, which are typically recalcitrant to receiving foreign DNA (10, 12, 17). Consequently, the process of cleavage-and-repairing, which is typically considered the important advantage of conventional CRISPR-Cas systems, becomes a bottleneck to perform CRISPR-Cas-based genome editing in acetogenic bacteria.

Lately, a new CRISPR-Cas-based genome-editing tool, namely base editing, was developed by combining a CRISPR-Cas system with a deamination system to achieve genome editing at a one-nucleotide resolution without the necessity for DNA cleavage, homology-directed repair, and donor DNA (18–21). By creating a fusion of a nuclease impaired Cas protein (*i.e.*, nuclease deactivated Cas9, dCas9) and a deaminase, this tool generates cytosine (C) to thymine (T) substitutions with cytidine deaminase (Fig. 1A), or adenine (A) to guanine (G) substitutions with adenosine deaminase (18, 22). Base editing provides distinctive advantages for genome editing in acetogenic bacteria by circumventing the bottlenecks of conventional CRISPR-Cas systems in bacteria. One such advantage is that the required DNA-uptake ability of acetogenic bacteria is minimized. Despite a few principal demonstrations in other bacteria (23–28), the potential of base editing in acetogenic bacteria has not yet been unraveled.

**Fig. 1.**
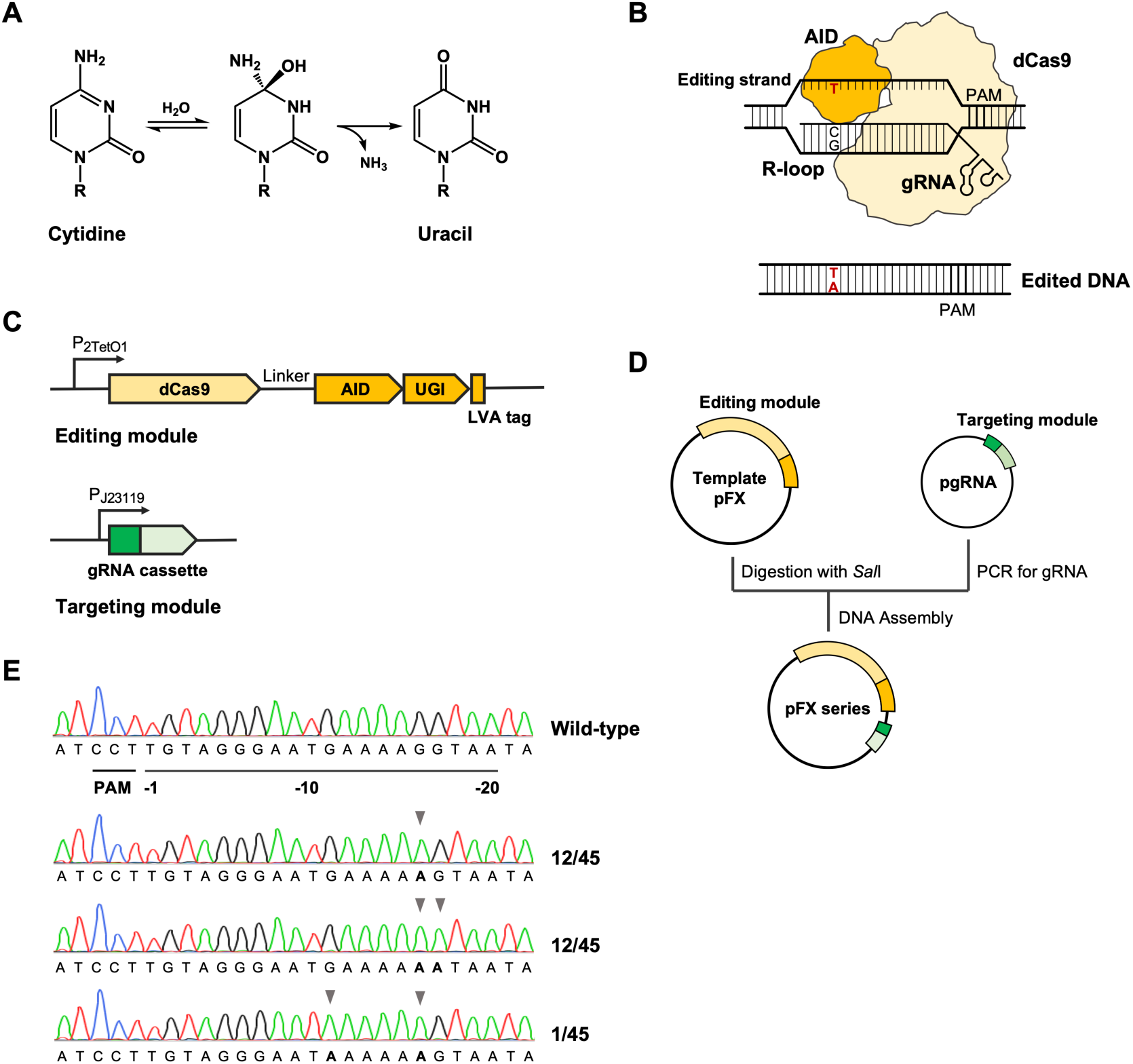
Design and validation of base editing in *C. ljungdahlii*. (A) Chemistry of deamination process converting cytidine to uracil. (B) Mechanism of base editing. Targeted by a gRNA, dCas9 binds to the target DNA and forms an R-loop. Activation-induced cytosine deaminase deaminates the Cs in the single strand DNA in the R-loop (Editing strand), resulting in C-to-T single-nucleotide variation in the genome. (C) The editing module consists of dCas9, activation-induced cytosine deaminase, uracil glycosylase inhibitor, and Leu-Val-Ala tag under the control of an inducible *tetR*-P*tet* system, and the targeting module contains the gRNA cassette under the control of the constitutive PJ23119 promoter. (D) Modularized strategy to generate editing plasmid series. To generate an editing plasmid, an inverse PCR is employed to generate the gRNA using pgRNA as a template. Template pFX plasmid is digested with *Sal*I and then assembled with the amplified gRNA cassette, resulting in the editing plasmid (pFX series). (E) Validation of base editing using *pta* as a target. The protospacer adjacent motif is displayed on the complimentary sequence, and the position of nucleotides in the protospacer were counted from the first nucleotide adjacent to the protospacer adjacent motif (position −1). Arrows indicate identified single-nucleotide variations. (PAM: protospacer adjacent motif; AID: activation-induced cytosine deaminase; UGI: uracil glycosylase inhibitor; and LVA tag: Leu-Val-Ala tag.)

Here, we developed a modularized base-editing tool for acetogenic bacteria by coupling dCas9 from *S. pyogenes* with activation-induced cytidine deaminase from the sea lamprey *Petromyzon marinus* (28). Efficient base editing was validated, and the editing principles were investigated in the model acetogenic bacterium *C. ljungdahlii*. Genome-scale *in-silico* analysis revealed the capability of our base-editing tool. As a first application, we employed base editing to reprogram the distribution of the carbon flux from acetyl-CoA to acetate and ethanol during heterotrophic and autotrophic fermentation, linking designed single-nucleotide variations with industrially relevant bacteria. Our base-editing tool will promote the research and application of C1 utilization with acetogenic bacteria, and more generally, provides an example for upgrading CRISPR-Cas-based genome-editing tools in bacteria.

## Results

### Design of a modularized base-editing tool for *C. ljungdahlii*

For our base-editing tool, we constructed a fusion of a dCas9 with an activation-induced cytidine deaminase. We selected the dCas9 (D10A and H840A) from *S. pyogenes* and combined it with the activation-induced cytidine deaminase from *P. marinus* to minimize the toxicity of Cas nuclease and to obtain promising deamination performance (28). The dCas9, together with a gRNA, serves as a navigator to target a specific DNA sequence (a protospacer), which has to be located next to a protospacer adjacent motif (PAM), on the genome. When binding to the target an R-loop is formed (Fig. 1B). Cytidine deaminase then converts cytidine to uracil *via* a deamination process (Fig. 1A). In the following replication or repair of the DNA, the cell reads uridine as T, which results in a C-to-T single nucleotide variation (28). The deamination occurs on the single-strand DNA (editing strand) in the R-Loop and changes C to T in a defined editing window (Fig. 1B) (18, 28). To increase the editing efficiency and to prevent the excision of uracil on the editing strand, an uracil glycosylase inhibitor was fused to an activation-induced cytidine deaminase. Furthermore, a fusion to a Leu-Val-Ala protein degradation tag was added, which leads to an overall lower amount of the fusion protein in the cell, to minimize the potential toxicity of dCas9 and uracil glycosylase inhibitor (28). Furthermore, we employed a tetracycline repressor-promoter (*tetR*-P_*tet*_) system, which is inducible with anhydrotetracycline in *C. ljungdahlii*, for the regulated expression of our base-editing tool (Fig. 1C) (29).

To introduce our base-editing tool to *C. ljungdahlii*, we designed a modularized plasmid system, which contains: **1**) a template plasmid (pFX) that carries the editing module, consisting of dCas9, activation-induced cytidine deaminase, uracil glycosylase inhibitor, and Leu-Val-Ala tag under the control of the inducible *tetR*-P_*tet*_ system; and **2**) a helper plasmid (pgRNA01) for the streamlined generation of the targeting modules (gRNAs, Fig. 1C). This modularization allows the use of inverse PCR on the helper plasmid to exchange the protospacer in the gRNA for a specific genome target site. The protospacer is driven by the P_J23119_ promoter and is flanked with the *S. pyogenes* Cas9 scaffold to form the new targeting module (*SI Appendix*, Fig. S1). Afterwards, the targeting module can be assembled with the pFX plasmid, creating the editing plasmid (Fig. 1D). Finally, *C. ljungdahlii* can be transformed with the methylated editing plasmids to mediate base editing.

### Validation of base editing in *C. ljungdahlii*

To validate our system, *pta* (CLJU_c12770) from *C. ljungdahlii*, which codes for the phosphotransacetylase, was selected as a first target (gRNA01, *SI Appendix*, Table S1). We discovered efficient conversion from C to T on the editing strand, leading to a G-to-A single-nucleotide variation in the coding strand (Fig. 1E). In total, 45 colonies from 5 individual rounds of base editing were picked to analyze the frequency and editing window. Three clean editing patterns were identified (Fig. 1E). The highest editing frequency was found for position −16 of the protospacer in all three patterns (counting the site adjacent to the protospacer adjacent motif as position −1). Twelve out of 45 colonies showed a single mutation at this position −16, 12 colonies showed double mutations at positions - 16 and −17, and 1 colony showed a double mutation at positions −12 and −16 (Fig. 1E). We also identified 2 colonies with mutations at position −11 (Fig. 1E) and −2 (*SI Appendix*, Fig. S2), respectively, outside of the editing window that was described by others (position −20 to −15) (18, 28), while the latter one (position −2) was a colony with mixed signals at position −17 (*SI Appendix*, Fig. S2). Importantly, this finding did not considerably influence the chance to select colonies with the desired single-nucleotide variations in only one round of selection. When targeting another site in *pta* (gRNA06, *SI Appendix*, Table S1), only 1 out of 8 colonies was found to be edited (*SI Appendix*, Fig. S3A), suggesting a sequence-specific editing preference, which is in agreement with previous reports (22).

We further interrogated the editing principles of the base-editing tool. Our base-editing tool converts C to T on the editing strand, which indicates that it only deaminates C(s) in the protospacer (18, 21, 28). Therefore, we designed two gRNAs with no Cs in the protospacers (gRNA02 and gRNA07, *SI Appendix*, Table S1) to examine the base-editing mechanism further, because we hypothesized that in this case no base editing would occur. Importantly, this experiment can hardly be done in non-A-T-rich bacteria in which protospacers without a C are much less abundant. As anticipated, we did not observe single-nucleotide variations in any colony (*SI Appendix*, Fig. S3B). Others have demonstrated that activation-induced cytidine deaminase-mediated base editing showed an editing window of five nucleotides starting from the opposite end of the protospacer adjacent motif, and that the window shifts depending on the length of protospacers (18, 21, 28). However, we discovered that our base-editing tool did not lead to mutations of the C at position −20 with neither a 20- or a 22-nucleotide protospacer (gRNA05 and gRNA15, *SI Appendix*, Table S1). This suggests an editing window in *C. ljungdahlii* starting from position −19 (*SI Appendix*, Fig. S3C). Additionally, we observed a wider editing window (position −19 to −2) within the protospacer, although the editing frequency was very low in the range between position −11 to −2 (Fig. 1E and *SI Appendix*, Fig. S2 and Fig. S3A).

### *In-silico* evaluation of base-editing capability on genome-scale

One doubt for applying base editing in acetogenic bacteria might be a questionable editing capability, because dCas9 from *S. pyogenes* recognizes the nucleotides NGG as the protospacer adjacent motif and *C. ljungdahlii* is an A-T-rich bacterium (77% A-T) (NCBI GenBank Access No. CP001666.1). To investigate the editing capability of our base-editing tool, we developed a genome-scale algorithm. The algorithm reads all possible protospacer adjacent motifs from the genome sequence, and then identifies Cs on the editing strand and converts those Cs to Ts within the editing window. This returns mutations at translational level and identifies the genome-wide capability to install missense mutations, nonsense mutations, and premature STOP codons (*Dataset*). To avoid an overestimation of the capability, we defined position −19 to −11 as the editing window, which combined our results and previous reports (18), although editing outside this window might still have been possible. We found that 314,800 sites could be potentially edited among which 257,133 sites are located in coding regions (Fig. 2A). These editable sites involve 99.83% (4,177 out of 4,184) of all genes. Only 7 genes that encode short hypothetic proteins (25 to 50 amino acids) cannot be edited (*SI Appendix*, Table S2). We found that: **1**) 99.69% (4,171 out of 4,184) genes can be edited to have missense mutations; **2**) 99.04% (4,144 out of 4,184) genes can be edited to have nonsense mutations; and **3**) 81.36% (3,404 out of 4,184) genes can be inactivated/truncated by installing premature STOP codons (30) (Fig. 2B, *SI Appendix*, Table S2). These *in-silico* results demonstrate a great capability of base editing even in an A-T-rich bacterium such as *C. ljungdahlii*.

**Fig. 2.**
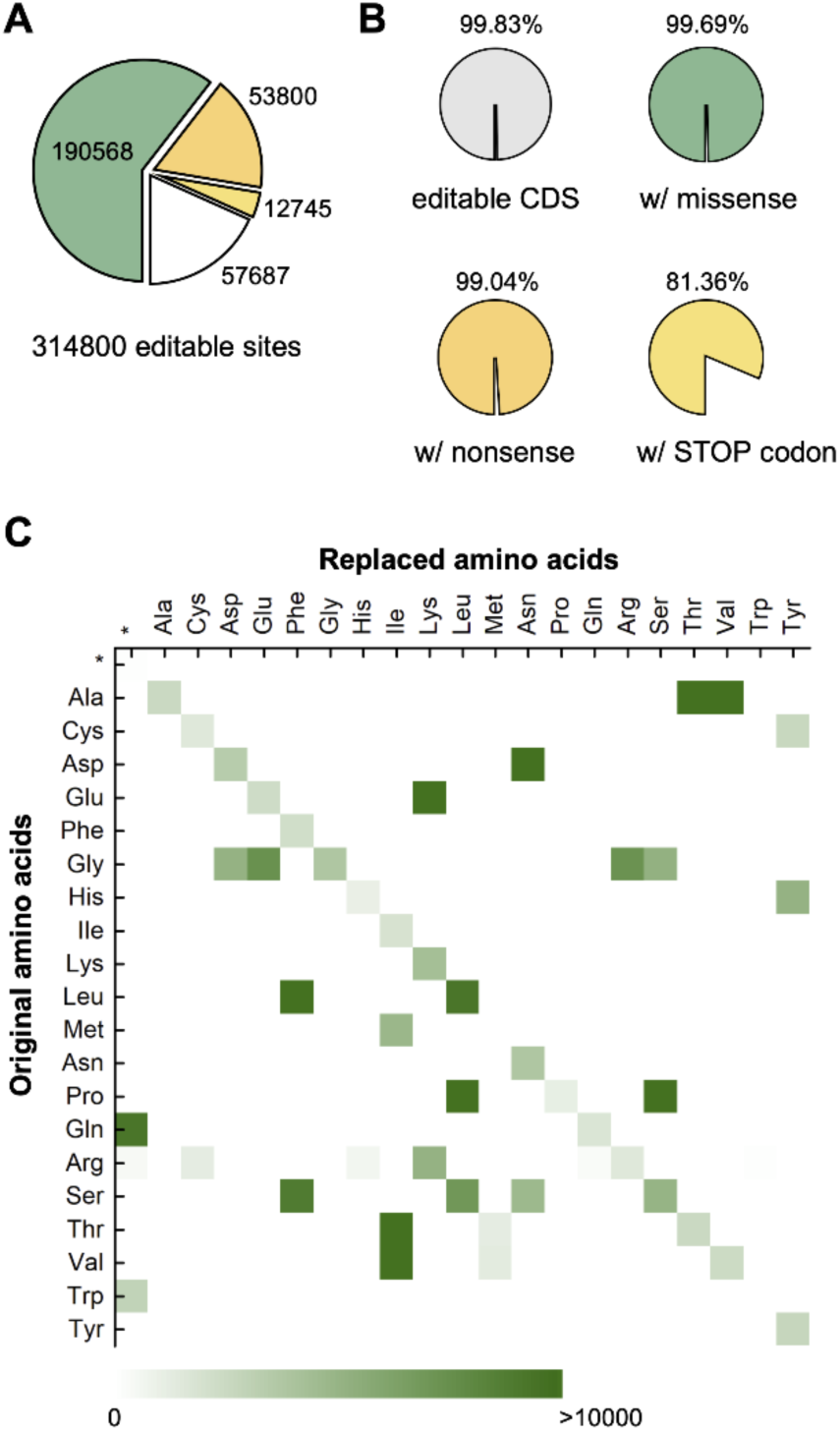
*In-silico* evaluation of base-editing capability in *C. ljungdahlii*. (A) Pie chart with the numbers of sites that can be edited on a genome scale. Green indicates missense mutations, orange indicates nonsense mutations (excluding premature STOP codons), yellow indicates premature STOP codons, and white indicates single-nucleotide variations that are not located in coding regions. (B) Pie charts with the percentages of genes that can be edited in generating different kinds of mutations in coding regions. (C) Amino acid replacement matrix generated by base editing. The green squares indicate the possible mutations with a lighter green color indicating fewer and a darker green color indicating more possible mutations on genome-scale.

On a genome scale, we discovered that, except for Trp and Met codons, all other amino acid codons can be edited to lead to nonsense mutations without changing the amino acid, while 15 out of 20 amino acids (excluding Phe, Ile, Lys, Asn, and Trp) can be changed to another amino acid (missense mutation) by changing the codon *via* single-nucleotide variations (Fig. 2C). Importantly, Gln, Arg, and Trp codons can be replaced to STOP codons. By changing CAA to TAA or CAG to TAG, 9,329 Gln codons can be changed to STOP codons. Arg (427 sites) also shows potential to be mutated to a STOP codon by converting CGA to TGA. Trp (2,989 sites) offers a different strategy to install premature STOP codons, because base editing changes CCA to TTA, TCA, or CTA on the editing strand and results in TAA, TGA, or TAG on the coding strand. Compared with a predicted amino acid replacement matrix in G-C-rich *Streptomyces* species, only 3 out of 32 different amino acid replacement routines were not identified for *C. ljungdahlii* (Pro to Phe, Gly to Lys, and Gly to Asn), which probably results from the A-T-rich genome (26) (Fig. 2C).

### Reprogramming carbon flux by installing premature STOP codons

To further demonstrate the application potential of our base-editing tool, we disrupted genes involved in ethanol production in *C. ljungdahlii* to reprogram the carbon flux for improved acetate production as a first application. To achieve this, we targeted four genes in two different metabolic pathways individually. First, we targeted *adhE1* (CLJU_c16510) and *adhE2* (CLJU_c16520), which encode isoenzymes of the bifunctional aldehyde-alcohol dehydrogenase. This enzyme converts acetyl-CoA to ethanol *via* acetaldehyde as an intermediate under heterotrophic conditions (Fig. 3A) (2, 8). The premature STOP codons were successfully installed in *adhE1* (with gRNA10, *SI Appendix*, Table S1) and *adhE2* (with gRNA11, *SI Appendix*, Table S1), respectively, generating strains QX3 (*adhE1* Trp169*) and QX4 (*adhE2* Gln33*) (Fig. 3B). Second, we targeted *aor1* (CLJU_c20110) and *aor2* (CLJU_c20210), which encode isoenzymes of the aldehyde:ferredoxin oxidoreductase. This enzyme converts acetate (in the form of undissociated acetic acid) to acetaldehyde under autotrophic conditions (Fig. 3A) (2). Accordingly, we inactivated *aor1* and *aor2* by installing STOP codons with gRNA19 and gRNA14 (*SI Appendix*, Table S1), generating strains QX5 (*aor1* Gln267*) and QX6 (*aor2* Gln267*), respectively (Fig. 3B).

**Fig. 3.**
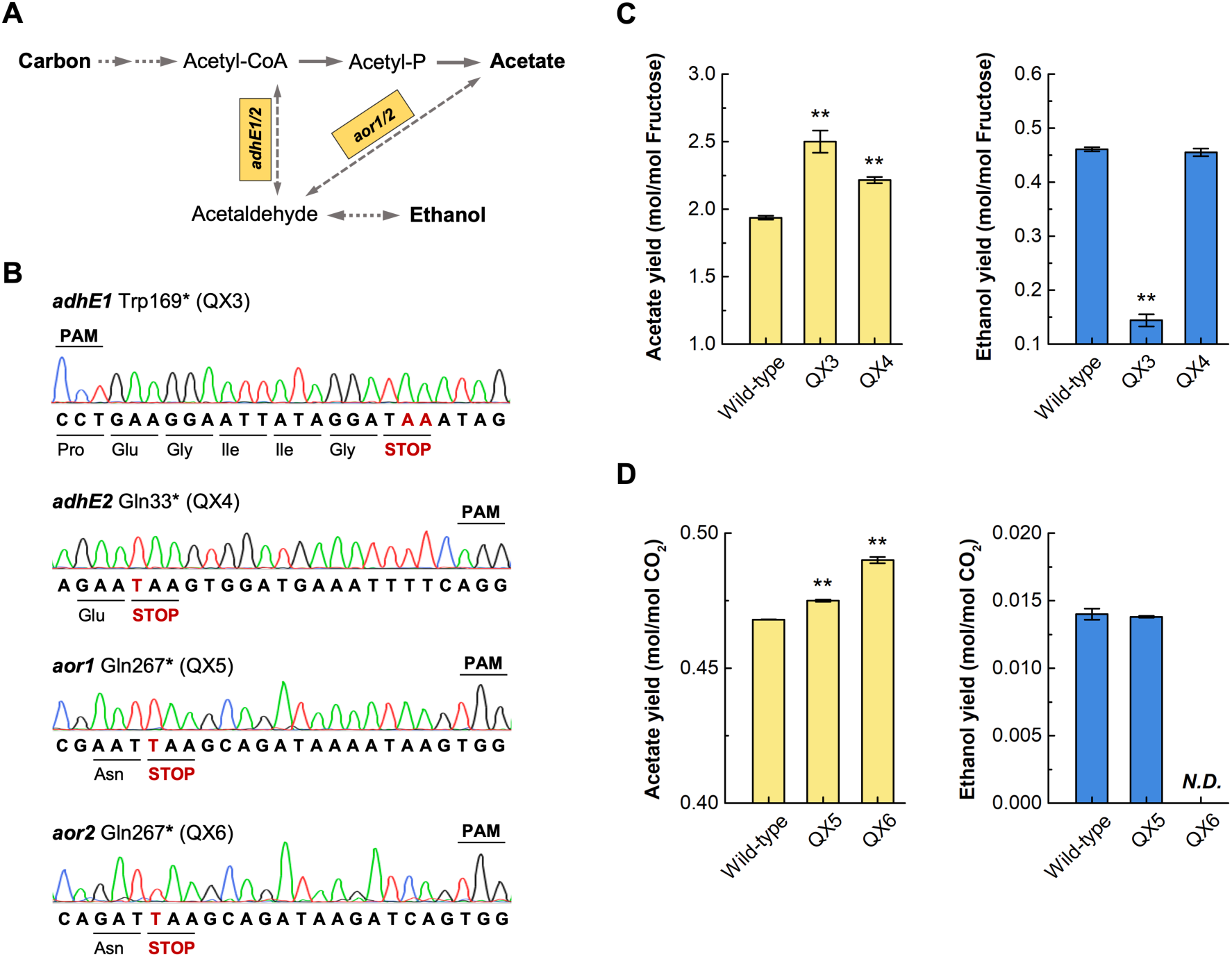
Reprogramming carbon flux towards improved acetate production. (A) Metabolic pathway from acetyl-CoA to acetate and ethanol. Under heterotrophic conditions, ADHE1/2 (encoded by *adhE1/2*) convert acetyl-CoA to ethanol *via* acetaldehyde as an intermediate, while under autotrophic conditions, AOR1/2 (encoded by *aor1/2*) convert acetate to acetaldehyde. (B) Validation of premature STOP codons in the four edited strains (QX3, QX4, QX5, and QX6). The edited sequences and amino acids are shown in the protospacer region. (C) Acetate and ethanol yields of wild-type, QX3, and QX4 under heterotrophic conditions with 5 g/L (27.8 mM) of fructose as the carbon source. (D) Acetate and ethanol yields of wild-type, QX5, and QX6 under autotrophic conditions with a gas mixture of H_2_/CO_2_ (80/20 vol-%, 1.5 bar) as the substrate. The fermentation experiments were conducted in triplicate (N=3), and the error bars indicate the standard deviations. The differences in acetate yield and ethanol yield were verified by *t*-test with a P < 0.05 as a significant difference and a P < 0.001 as a highly significant difference (**).

We first tested the stability of all four strains by serial transfer experiments. We confirmed the correct genotype (single nucleotide variations) at the edited location after 10 transfers with more than 65 generations, indicating that our base-editing tool resulted in stable genotypes (*SI Appendix*, Fig. S4). Next, we investigated the physiology with bottle experiments. For heterotrophic conditions, QX3 and QX4 showed growth defects compared to the wild-type strain, and did not consume all of the provided fructose (*SI Appendix*, Fig. S5A and S5B). However, both QX3 and QX4 achieved higher final acetate yields, because at similar final biomass and acetate concentrations, these strains achieved lower final ethanol concentrations (Fig. 3C, *SI Appendix*, Fig. S5C and S5D). Especially for QX3, we observed a 28.9% higher acetate yield and a 68.6% reduced ethanol yield compared to the wild-type strain (Fig. 3C), which is in agreement with a previous report on an *adhE1* deletion in *C. ljungdahlii* (8). For autotrophic conditions, we also found growth defects for QX5 and QX6 compared to the wild-type strain, and less overall substrate (H_2_/CO_2_) consumption (*SI Appendix*, Fig. S6A and S6B). QX5 showed wild-type-like patterns in the yield of acetate and ethanol, however, at an overall lower absolute level of final concentrations (Fig. 3D, *SI Appendix*, Fig. S6C and S6D). QX6 showed a slight, but significant, increase of 4.6% in the final acetate yield (0.490 mol acetate/mol consumed CO_2_) compared to the wild-type strain (0.468 mol acetate/mol consumed CO_2_), while ethanol was below our detection limit (Fig. 3D, *SI Appendix*, Fig. S6C). The higher acetate yield likely resulted from a redistribution of carbon from biomass and/or ethanol to acetate production. The final biomass and acetate concentrations were considerably lower compared to the wild-type strain (*SI Appendix*, Fig. S6A and S6D), while we further acknowledge that this increase (4.6%) in acetate yield by QX6, although significant, is only marginal. However, this increase brings the acetate yield (0.490 mol acetate/mol consumed CO_2_) closer to the theoretical limit (0.500 mol acetate/mol consumed CO_2_). We found that a single-gene inactivation (*adhE1* and *aor2*) could be enough to generate strains with higher acetate yield and lower ethanol yield under either heterotrophic or autotrophic conditions. To further optimize the metabolism, it is also possible to obtain multigene inactivation in one strain with multiplex base editing by employing established assembly approaches to generate gRNA arrays with protospacers targeting different genes as described by others (28, 29).

## Discussion

### An expanded synthetic biology toolkit for acetogenic bacteria

We developed a base-editing tool for the model acetogenic bacterium *C. ljungdahlii* and enabled genome editing at a one-nucleotide resolution without DNA cleavage, homology-directed repair, and donor DNA. Base editing bypasses the general bottlenecks of applying CRISPR-Cas systems in bacteria, which include the toxicity of Cas nucleases and inefficient DNA repairing mechanisms. It also lowers the requirement of transformation efficiency in *C. ljungdahlii* compared to conventional CRISPR-Cas-based genome editing (10, 12, 16). We discovered a great capability of our base-editing tool. With base editing, we can: **1**) install STOP codons to 3,404 genes in *C. ljungdahlii* to reprogram the metabolisms directly; **2**) generate nonsense mutations in 4,144 genes to interrogate codon preference; and **3**) replace amino acids *via* missense mutations in 4,171 genes to perform protein research and engineering *in vivo*. Moreover, we observed that desired single-nucleotide variations could be obtained in a single round of selection. Only in two cases, we observed both wild-type and edited signals in one colony. In a previous study, mixed populations have been reported as an issue for base editing in *C. berjerinkii*, making a second round of selection necessary (27). Notably, we observed high precision single-nucleotide variations in *C. ljungdahlii* with limited bystander nucleotide substitutions (undesired single-nucleotide variations within the editing window on the editing strand) (Fig. 3B). This is an advantage returned by the A-T-rich genome of *C. ljungdahlii*, which naturally overcomes bystander base editing with limited Cs in a target sequence and leads to precise base editing. We designed a modularized system to enable fast generation of the base-editing plasmid series. The employed plasmid backbone, replicon for clostridia, antibiotic resistances markers, and the dCas9 protein have been separately demonstrated to be functional in various species in the order *Clostridiales*, including *Acetobacterium woodii* (dCas9 has not yet been validated) (31, 32), *Eubacterium limosum* (13), and *Clostridium autoethanogenum* (14). Accordingly, the system could be easily generalized in acetogenic bacteria, which mainly belong to the order *Clostridiales*.

### Linking base editing with microbial C1 utilization

Base editing was first invented to revert single-nucleotide mutations related to human diseases (20). Despite an increasing utilization in medicine and agriculture (18, 22), only a few reports validated bacterial base-editing principles, especially for bacterial pathogens (23–25). Furthermore, only few reports exist for biotechnologically relevant bacteria, and these do not demonstrate a specific biotechnological application (26–28). Presumably, base editing in bacteria might be hindered by a low editing capability in relevant genes or low stability of resulting single-nucleotide variations, which would not be favorable for industrial biotechnology. To overcome this presumption, we inactivated four genes related to ethanol production in *C. ljungdahlii* as a first application, with the goal to increase the acetate yield. This would improve the production of certain platform chemicals that require acetate as an intermediate (33, 34). For instance, our acetate-producing strain can be considered to further improve the two-stage bioprocess for single-cell protein production from C1 gases, with acetate as the carbon-fixing intermediate product before being fed to aerobic yeasts, especially for industrial gases that contain CO (4). Importantly, single-nucleotide variations generated by base editing are clean mutations in the genome, which may also occur in natural evolution. Thus, our base-editing tool, in principle, provides a unique venue to engineer industrially relevant bacteria without creating genetically modified organisms (GMOs). However, this advantage is not recognized in the legislation of all countries, and especially in Europe, CRISPR-Cas-based genome editing is often *per se* considered to generate GMOs, irrespective of the outcome of the editing.

### Limitations and perspectives for base editing in A-T-rich bacteria

Not surprisingly, base editing has its limitations in A-T-rich bacteria. First, the editing sites are still limited in A-T-rich genomes, because of the protospacer adjacent motif (nucleotides NGG) that is recognized by dCas9 from *S. pyogenes*. Despite a large number of editable sites, not the entire genome can be covered. To overcome this, possible strategies include using a dCas protein with a different protospacer adjacent motif such as Cas12a with nucleotides TTTV as a protospacer adjacent motif (35) or xCas9 with nucleotides NG as a protospacer adjacent motif (36). Second, base editing is site-specific, and not all sites following the editing principles can be edited (22). Evidently, when we tried to introduce a STOP codon at Gln235 in *aor1* with gRNA13, no colonies with the expected single-nucleotide variations were identified (data not shown), while we obtained 2 out of 8 colonies with Gln237* replacement by using gRNA19. A different deaminase may be necessary to circumvent this limitation. Third, base editing intrinsically cannot insert DNA fragments into the genome. Yet, it offers a new angle to edit the genome with CRISPR-Cas systems without DNA cleavage. Starting from this perspective, a recent report demonstrated DNA insertion into the genome without cutting the DNA by coupling a CRISPR-Cas system to a reverse-transcriptase (37).

In summary, we established an efficient base-editing tool for gene manipulation in acetogenic bacteria. Further, we demonstrated the use of this cutting-edge genome-editing tool in C1 utilization with the industrially relevant acetogenic bacterium *C. ljungdahlii*. Our strategy provides an example for upgrading bacterial genome-editing tools with CRISPR systems in general, especially for bacteria that are sensitive to heterologously expressed Cas nucleases (*e.g.*, cyanobacteria) and those with limited capability of receiving foreign DNA.

## Methods and Materials

Methods and Materials for: strains and media, plasmid constructions, transformation of *C. ljungdahlii*, base editing in *C. ljungdahlii*, plasmid curing, serial transfer experiments, fermentation experiments, and genome-scale algorithm design are described in *SI Appendix*.

## Acknowledgement

This work was supported by the Alexander von Humboldt Foundation in the framework of the Alexander von Humboldt Professorship (L.T.A.) and the Humboldt Research Fellowship for Postdoctoral Researchers (P.-F.X.). S.S. thanks the support from the German Federal Environmental Foundation (Deutsche Bundesstiftung Umwelt, DBU). B.M. and L.T.A are grateful to funding from the Deutsche Forschungsgemeinschaft (DFG, German Research Foundation) under Germany’s Excellence Strategy – EXC 2124 – 390838134. Finally, LTA is supported as a Max Planck Fellow by the Max Planck Institute for Developmental Biology.

## Author contributions

P.-F.X. designed the experiments. P.-F.X., I.C., S.S., and C.K. performed research, and analyzed data. B.M. and L.T.A supervised the project. P.-F.X. and B.M. wrote the paper, while all edited the manuscript.

## References

1. M. Köpke et al., *Clostridium ljungdahlii* represents a microbial production platform based on syngas. Proc. Natl. Acad. Sci. U. S. A. 107, 13087–13092 (2010).

2. H. Richter et al., Ethanol production in syngas-fermenting *Clostridium ljungdahlii* is controlled by thermodynamics rather than by enzyme expression. Energy Environ. Sci. 9, 2392–2399 (2016).

3. T. Ueki, K. P. Nevin, T. L. Woodard, D. R. Lovley, Converting carbon dioxide to butyrate with an engineered strain of *Clostridium ljungdahlii*. mBio 5, e01636–01614 (2014).

4. B. Molitor, A. Mishra, L. T. Angenent, Power-to-protein: converting renewable electric power and carbon dioxide into single cell protein with a two-stage bioprocess. Energy Environ. Sci. 12, 3515–3521 (2019).

5. B. Molitor et al., Carbon recovery by fermentation of CO-rich off gases - Turning steel mills into biorefineries. Bioresour. Technol. 215, 386–396 (2016).

6. V. Müller, New horizons in acetogenic conversion of one-carbon substrates and biological hydrogen storage. Trends Biotechnol. 37, 1344–1354 (2019).

7. A. Banerjee, C. Leang, T. Ueki, K. P. Nevin, D. R. Lovley, Lactose-inducible system for metabolic engineering of *Clostridium ljungdahlii*. Appl. Environ. Microbiol. 80, 2410–2416 (2014).

8. C. Leang, T. Ueki, K. P. Nevin, D. R. Lovley, A genetic system for *Clostridium ljungdahlii*: a chassis for autotrophic production of biocommodities and a model homoacetogen. Appl. Environ. Microbiol. 79, 1102–1109 (2013).

9. G. J. Knott, J. A. Doudna, CRISPR-Cas guides the future of genetic engineering. Science 361, 866–869 (2018).

10. H. Huang et al., CRISPR/Cas9-based efficient genome editing in *Clostridium ljungdahlii*, an autotrophic gas-fermenting bacterium. ACS Synth. Biol. 5, 1355–1361 (2016).

11. H. Huang, C. Chai, S. Yang, W. Jiang, Y. Gu, Phage serine integrase-mediated genome engineering for efficient expression of chemical biosynthetic pathway in gas-fermenting *Clostridium ljungdahlii*. Metab. Eng. 52, 293–302 (2019).

12. R. Zhao et al., CRISPR-Cas12a-mediated gene deletion and regulation in *Clostridium ljungdahlii* and its application in carbon flux redirection in synthesis gas fermentation. ACS Synth. Biol. 8, 2270–2279 (2019).

13. J. Shin et al., Genome engineering of *Eubacterium limosum* using expanded genetic tools and the CRISPR-Cas9 system. ACS Synth. Biol. 8, 2059–2068 (2019).

14. S. Nagaraju, N. K. Davies, D. J. F. Walker, M. Köpke, S. D. Simpson, Genome editing of *Clostridium autoethanogenum* using CRISPR/Cas9. Biotechnol. Biofuels 9, 219 (2016).

15. K. Selle, R. Barrangou, Harnessing CRISPR-Cas systems for bacterial genome editing. Trends Microbiol. 23, 225–232 (2015).

16. J. M. Vento, N. Crook, C. L. Beisel, Barriers to genome editing with CRISPR in bacteria. J. Ind. Microbiol. Biotechnol. 46, 1327–1341 (2019).

17. B. Molitor, K. Kirchner, A. W. Henrich, S. Schmitz, M. A. Rosenbaum, Expanding the molecular toolkit for the homoacetogen *Clostridium ljungdahlii*. Sci. Rep. 6, 31518 (2016).

18. H. A. Rees, D. R. Liu, Base editing: precision chemistry on the genome and transcriptome of living cells. Nat. Rev. Genet. 19, 770–788 (2018).

19. N. M. Gaudelli et al., Programmable base editing of A•T to G•C in genomic DNA without DNA cleavage. Nature 551, 464–471 (2017).

20. A. C. Komor, Y. B. Kim, M. S. Packer, J. A. Zuris, D. R. Liu, Programmable editing of a target base in genomic DNA without double-stranded DNA cleavage. Nature 533, 420–424 (2016).

21. K. Nishida et al., Targeted nucleotide editing using hybrid prokaryotic and vertebrate adaptive immune systems. Science 353, aaf8729 (2016).

22. K. A. Molla, Y. Yang, CRISPR/Cas-mediated base editing: technical considerations and practical applications. Trends Biotechnol. 37, 1121–1142 (2019).

23. Y. Wang et al., CRISPR-Cas9 and CRISPR-assisted cytidine deaminase enable precise and efficient genome editing in *Klebsiella pneumoniae*. Appl. Environ. Microbiol. 84, e01834–01818 (2018).

24. W. Chen et al., CRISPR/Cas9-based genome editing in *Pseudomonas aeruginosa* and cytidine deaminase-mediated base editing in *Pseudomonas* species. iScience 6, 222–231 (2018).

25. T. Gu et al., Highly efficient base editing in *Staphylococcus aureus* using an engineered CRISPR RNA-guided cytidine deaminase. Chem. Sci. 9, 3248–3253 (2018).

26. Y. Tong et al., Highly efficient DSB-free base editing for streptomycetes with CRISPR-BEST. Proc. Natl. Acad. Sci. U. S. A. 116, 20366–20375 (2019).

27. Q. Li et al., CRISPR-Cas9(D10A) nickase-assisted base editing in the solvent producer *Clostridium beijerinckii*. Biotechnol. Bioeng. 116, 1475–1483 (2019).

28. S. Banno, K. Nishida, T. Arazoe, H. Mitsunobu, A. Kondo, Deaminase-mediated multiplex genome editing in *Escherichia coli*. Nat. Microbiol. 3, 423–429 (2018).

29. B. M. Woolston, D. F. Emerson, D. H. Currie, G. Stephanopoulos, Rediverting carbon flux in *Clostridium ljungdahlii* using CRISPR interference (CRISPRi). Metab. Eng. 48, 243–253 (2018).

30. C. Kuscu et al., CRISPR-STOP: gene silencing through base-editing-induced nonsense mutations. Nat. Methods 14, 710–712 (2017).

31. M. H. Beck, M. Flaiz, F. R. Bengelsdorf, P. Dürre, Induced heterologous expression of the arginine deiminase pathway promotes growth advantages in the strict anaerobe *Acetobacterium woodii*. Appl. Microbiol. Biotechnol. 104, 687–699 (2020).

32. S. Hoffmeister et al., Acetone production with metabolically engineered strains of *Acetobacterium woodii*. Metab. Eng. 36, 37–47 (2016).

33. K. P. Nevin et al., Electrosynthesis of organic compounds from carbon dioxide is catalyzed by a diversity of acetogenic microorganisms. Appl. Environ. Microbiol. 77, 2882–2886 (2011).

34. S. N. Nangle, K. K. Sakimoto, P. A. Silver, D. G. Nocera, Biological-inorganic hybrid systems as a generalized platform for chemical production. Curr. Opin. Chem. Biol. 41, 107–113 (2017).

35. X. Li et al., Base editing with a Cpf1-cytidine deaminase fusion. Nat. Biotechnol. 36, 324–327 (2018).

36. J. H. Hu et al., Evolved Cas9 variants with broad PAM compatibility and high DNA specificity. Nature 556, 57–63 (2018).

37. A. V. Anzalone et al., Search-and-replace genome editing without double-strand breaks or donor DNA. Nature 576, 149–157 (2019).

