## Supplementary material for "Reprogramming acetogenic bacteria with CRISPR-targeted base editing *via* deamination": SI Appendix

### Methods and Materials

#### Strains and Media

*Clostridium ljungdahlii* DSM13528 was used as the wild-type strain, and all *C. ljungdahlii* strains that were used in this study are listed in Table S3. Reinforced clostridial medium (RCM) was employed for general cultivation of *C. ljungdahlii* (1). Modified PETC medium (ATCC 1754 medium) was used for fermentation experiments with 2 g/L (10 mM) of 2-(N-morpholino)ethanesulfonic acid buffering the medium instead of $NaHCO_3$. For heterotrophic fermentation, 5 g/L (27.8 mM) of fructose were added in 250 mL serum bottles with 100 mL of medium as the carbon source. For autotrophic fermentation, an $H_2/CO_2$ mixture (80/20 vol-%, 1.5 bar) was added to the headspace of 1 L cultivation bottles (Pressure plus Duran bottle, VWR) with 100 mL of medium. The headspace was refilled to 1.5 bar every 24 h during fermentation. Clarithromycin (5 μg/L) was added when necessary. The manipulation of *C. ljungdahlii* was performed in an anaerobic chamber (UNILab Pro Eco, MBraun, Germany) with an $O_2$ level below 10 ppm. The cultivation was performed at 37°C. *E. coli* TOP10 (Invitrogen) was used for general cloning and gene manipulation. *E. coli* was cultivated in Luria-Bertani (LB) medium. Ampicillin (100 μg/L), spectinomycin (60 μg/L), and erythromycin (400 μg/L on plates and 250 μg/L for liquid medium) were used to select and maintain plasmids in *E. coli*.

#### Plasmid construction

The plasmids used in this study are summarized in Table S4. pMTLdCas9 was a generous gift from Gregory Stephanopoulos (2), pTargetF (Addgene plasmid # 62226) was a gift from Sheng Yang (3), and pScI_dCas-CDA-UL (Addgene plasmid # 108551) was a gift from Akihiko Kondo (4). The pgRNA01 plasmid was generated *via* inverse PCR using pTargetF as a template to recover the original $P_{J23119}$ promoter and create the gRNA01 cassette, generating pgRNA01 (Fig. S1). Then, the pgRNA plasmid series containing the designed gRNA cassettes were constructed *via* inverse PCR using pgRNA01 as a template. The pFX template was constructed by flanking the fragment consisting of activation-induced cytidine deaminase, uracil glycosylase inhibitor, and Leu-Val-Ala tag from pScI_dCas-CDA-UL with dCas9 on pMTLdCas9. Based on the pFX and pgRNA plasmid series, a modularized method was employed to construct the pFX plasmid series for base editing. First, the pFX template was digested with *Sal*I and the gRNA cassette was amplified by primers EBT-PFX-88 and -89. Second, the pFX plasmid series was generated by combining these two parts *via* Gibson assembly (New England labs, NEB) (Fig. 1D). Plasmids were methylated *via* co-transformation with pANA1, which carries Φ3tI methyltransferase before transformation of *C. ljungdahlii* with the respective plasmid (1). PCRs were performed using Q5 polymerase (NEB). The primers (Integrated DNA Technologies, BVBA, Leuven, Belgium) for plasmid construction are summarized in Table S5.

**Transformation of *C. ljungdahlii***

Transformation of *C. ljungdahlii* was following a modified protocol previously established (1). Briefly, RCM was inoculated and transferred twice with *C. ljungdahlii* at a 1:100 dilution. When the $OD_{600}$ in the second culture reached 0.2 - 0.4, 12 - 16 mL of culture were harvested by stepwise centrifugation at 10,000 rpm for 1.5 min (mySPIN™ 12 Centrifuge, Thermo Fisher Scientific). The cells were then washed with ice-cold glycerol (10 vol-%) for three times and resuspended in 200 μL of 10 vol-% glycerol as electrocompetent cells. The methylated plasmids (2 μg) were mixed with the competent cells individually for electroporation (Micropulse system, 2.5 kV, 600 Ω and 25 μF, Bio-Rad) in 2-mm cuvettes. After electroporation, the cells were immediately transferred to pre-warmed RCM for a 12 - 18 h recovery. Subsequently, RCM medium with 5 μg/L of clarithromycin was inoculated with the cells, and the outgrowth cultures were used directly for base editing.

**Base editing in *C. ljungdahlii***

The methylated pFX plasmid was transformed into *C. ljungdahlii* according to the protocol above. After electroporation, fresh RCM with 5 µg/L of clarithromycin and 100 ng/mL of anhydrotetracycline was inoculated (1:10 dilution) with the outgrowth culture (RCM with only clarithromycin) in a Hungate-type anaerobic culture tube and incubated for 18 - 24 h to induce base editing. After the induction, the culture was mixed with molten RCM agar (1.0 weight-%) with clarithromycin and poured into petri dishes. Single colonies were picked for further analysis. First, a colony PCR was conducted using Phire Plant Direct PCR Master Mix (Thermo Fisher Scientific) to amplify the fragment containing the target DNA sequence. Second, the fragment was sequenced to verify the single-nucleotide variations. Primers for amplifying the edited DNA fragments and sequencing are listed in Table S6.

**Plasmid curing**

To cure the cells from the plasmid, a colony with the designed single-nucleotide variations was used to re-inoculate fresh RCM without clarithromycin and transferred for a second time. The second culture was poured with RCM agar (1.0 weight-%) at different dilutions. Subsequently, single colonies were picked to determine the loss of plasmids using colony PCR. If the colony PCR showed no PCR signal, the colony was further tested in RCM and RCM with clarithromycin to identify the curation from the plasmid. A colony that: **1)** carries the designed single-nucleotide variations; **2)** has no PCR signal for the editing plasmid; and **3)** fails to grow in selective medium, was regarded as an edited strain for subsequent stability evaluation and fermentation experiments.

**Serial transfer experiments**

To test the stability of single-nucleotide variations in the edited strains, the obtained QX3, QX4, QX5, and QX6 strains were used to inoculate RCM from single colonies and transferred into fresh RCM at a 1:100 dilution after the $OD_{600}$ reached late exponential or stationary phase ($OD_{600}$ 1.3 to 2.0). The single-nucleotide variations were tested after each transfer and the sequencing results after the 10th transfer are shown as a demonstration of the stability. The number of generations (n) was calculated by the following equation:

$$n = \frac{\log(OD_t) - \log(OD_0)}{\log 2}$$

where $OD_t$ is the $OD_{600}$ before each transfer and $OD_0$ is the initial $OD_{600}$ after each transfer. The total number of generations is the sum of the generation numbers of each transfer.

**Fermentation experiments**

Fresh PETC medium was inoculated with *C. ljungdahlii* strains (from RCM cultures) and adapted to this medium in two transfers. Cells from the second preculture were harvested at late exponential phase and washed before inoculation. For heterotrophic fermentation, the wild-type, QX3, and QX4 strains were tested, and for autotrophic fermentation, the wild-type, QX5, and QX6 strains were tested. For all experiments, the initial $OD_{600}$ was adjusted to 0.1 and samples were taken at different time intervals to analyze the growth and products. The fermentation products, including acetate, ethanol, and the substrate fructose, were measured using HPLC (LC20, Shimazu, Japan) with a RID detector and Aminex HPX-87H column (oven temperature 65°C) using 5 mM $H_2SO_4$ as elution solvent (0.6 mL/min).

The acetate and ethanol yields in the heterotrophic fermentations were calculated as following:

$$\text{Yield (mol/mol)} = \frac{C_{Acetate/Ethanol,t}}{C_{Fructose,0} - C_{Fructose,t}}$$

Where $C_{Acetate/Ethanol,t}$ is the concentration of acetate or ethanol at 107.50 h; $C_{Fructose,0}$ is the concentration of fructose at time 0; and $C_{Fructose,t}$ is the concentration of fructose at 107.50 h.

For autotrophic fermentation, the consumed $CO_2$ was first calculated by adding up the carbon in biomass ($C_5H_8O_2N$), acetate, and ethanol. 100 mL of a *C. ljungdahlii* culture with an $OD_{600}$ of 1.0 equals 24.4 mg of biomass in dry cell weight (5), which corresponds to 0.214 mmol of carbon. Then the yield was calculated by:

$$\text{Yield (mol/mol)} = \frac{C_{Acetate/Ethanol,t}}{\text{consumed } CO_2 \text{ per culture volume}}$$

Where $C_{Acetate/Ethanol,t}$ is the concentration of acetate or ethanol at 137.75 h.

**Genome-scale algorithm design**

We developed an algorithm to identify all editable sites at genome-scale to identify possible single-nucleotide variations and mutations at translation level. The algorithm was coded in Python (Pycharm 2018.1 with a virtual environment), and the commented scripts and necessary files have been uploaded to GitHub (https://github.com/isacasini/SNV_Xia_et_al_2020). In brief, the algorithm first reads both strands of the genomic DNA of *C. ljungdahlii* (NCBI GenBank, Access No. CP001666.1), as well as an additional file with the necessary genomic information (*i.e.*, NCBI identifier, NCBI annotations, *etc.*), and it identifies all protospacers by locating the protospacer adjacent motifs (nucleotides NGG) as potential editable sites. According to the editing principles of our base editing system, the algorithm finds Cs in the identified protospacers between position -19 and -11. If a C (or several Cs) is found, it considers the site as an editable site and changes C to T, returning the single-nucleotide variations. If the editable site is located in a coding region, the algorithm also returns the resulting changes in amino acid. All editable sites and analysis were summarized in the *Dataset* and Table S2.

**Table S1** gRNA and protospacer adjacent motifs (PAM) sequences used in this study

| gRNA | Targets[a] | Strand[b] | PAM | gRNA sequence |
|---|---|---|---|---|
| gRNA01 | *pta* | N | AGG | TTACCTTTTCATTCCCTACA |
| gRNA02 | *pta* | C | AGG | AAAAATTTGGAGTAAGGCAA |
| gRNA04 | *pta* | C | AGG | GAAAAAATTTGGAGTAAGGCAA |
| gRNA05 | *pta* | C | AGG | CAAATAGTAAAGACAGCTCC |
| gRNA06 | *pta* | N | TGG | AATTGATCACTATCTGGGCA |
| gRNA10 | *adhE1* | N | AGG | ATCCATCCTATAATTCCTTC |
| gRNA11 | *adhE2* | C | AGG | AACAAGTGGATGAAATTTTC |
| gRNA13 | *aor1* | C | TGG | GGTCAGGGAATGCCAACTTA |
| gRNA14 | *aor2* | C | TGG | GATCAAGCAGATAAGATCAG |
| gRNA15 | *pta* | C | AGG | TTCAAATAGTAAAGACAGCTCC |
| gRNA19 | *aor1* | C | TGG | AATCAAGCAGATAAAATAAG |

[a] *pta* (CLJU_c12770), *adhE1* (CLJU_c16510), *adhE2* (CLJU_c16520), *aor1* (CLJU_c20110), and *aor2* (CLJU_c20210).

[b] N indicates non-coding strand, and C indicates coding strand.

**Table S2** Summary of the *in-silico* evaluation of our base editing tool

| Editing Type | Editable sites | Editable genes |
|---|---|---|
| Missense mutation | 190568 | 4171 |
| Nonsense mutation[a] | 66545 | 4144 |
| Premature STOP codon | 12745 | 3404 |
| Non-editable[b] | - | 7 |

[a] The number of nonsense mutations includes the number of premature STOP codons.
[b] Non-editable genes include: CLUJ_c02010 (44 amino acids); CLJU_c04140 (25 amino acids); CLJU_05590 (35 amino acids); CLJU_c06880 (40 amino acids); CLJU_c08940 (32 amino acids); CLJU_c18840 (43 amino acids); and CLJU_c35990 (50 amino acids).

**Table S3** *Clostridium ljungdahlii* strains used in this study

| **Strains** | **Description** | **References** |
|---|---|---|
| DSM13528 | *Clostridium ljungdahlii* type strain | DSM13528 |
| QX3 | DSM13528, *adhE1* Trp169* | This study |
| QX4 | DSM13528, *adhE2* Gln33* | This study |
| QX5 | DSM13528, *aor1* Gln267* | This study |
| QX6 | DSM13528, *aor2* Gln267* | This study |

**Table S4** Plasmids used in this study

| Name | Description | References |
|---|---|---|
| pANA1 | p15A, *bla*, Φ3tI methyltransferase | (1) |
| pMTLdSpCas9 | pMTL82254, $P_{2tetO1}$:*dcas9* | (2) |
| pTargetF | ColE1, *smR* | (3) |
| pScI_dCas9-CDA-UL | pSC101, *bla*, *dcas9*-PmCDA1-UGI-LVA | (4) |
| pgRNA01 | pTargetF, $P_{J23119}$:gRNA01 | This study |
| pgRNA02 | pgRNA01, $P_{J23119}$:gRNA02 | This study |
| pgRNA04 | pgRNA01, $P_{J23119}$:gRNA04 | This study |
| pgRNA05 | pgRNA01, $P_{J23119}$:gRNA05 | This study |
| pgRNA06 | pgRNA01, $P_{J23119}$:gRNA06 | This study |
| pgRNA10 | pgRNA01, $P_{J23119}$:gRNA10 | This study |
| pgRNA11 | pgRNA01, $P_{J23119}$:gRNA11 | This study |
| pgRNA13 | pgRNA01, $P_{J23119}$:gRNA13 | This study |
| pgRNA14 | pgRNA01, $P_{J23119}$:gRNA14 | This study |
| pgRNA15 | pgRNA01, $P_{J23119}$:gRNA15 | This study |
| pgRNA19 | pgRNA01, $P_{J23119}$:gRNA19 | This study |
| pFX | pMTLdSpCas9, dcas9 fused with PmCDA-UGI-LVA tag | This study |
| pFX01 | pFX, $P_{J23119}$:gRNA01 | This study |
| pFX02 | pFX, $P_{J23119}$:gRNA02 | This study |
| pFX04 | pFX, $P_{J23119}$:gRNA04 | This study |
| pFX05 | pFX, $P_{J23119}$:gRNA05 | This study |
| pFX06 | pFX, $P_{J23119}$:gRNA06 | This study |
| pFX10 | pFX, $P_{J23119}$:gRNA10 | This study |
| pFX11 | pFX, $P_{J23119}$:gRNA11 | This study |
| pFX13 | pFX, $P_{J23119}$:gRNA13 | This study |
| pFX14 | pFX, $P_{J23119}$:gRNA14 | This study |
| pFX15 | pFX, $P_{J23119}$:gRNA15 | This study |
| pFX19 | pFX, $P_{J23119}$:gRNA19 | This study |

**Table S5** Primers used for plasmid construction

| Primer | Sequence | Note |
|---|---|---|
| EBT-PFX-080 | GATTTGAGTCAGCTAGGAGGTGACGGTGGAGGAGGTTCTGGAGG | pFX series |
| EBT-PFX-086 | ATGCCTGGAGATCCTTACTCGAGTTATGCAACCAGTCCTAGCATCTTG | |
| EBT-PFX-081 | CCTCCAGAACCTCCTCCACCGTCACCTCCTAGCTGACTCAAATC | |
| EBT-PFX-087 | CAAGATGCTAGGACTGGTTGCATAACTCGAGTAAGGATCTCCAGGCAT | |
| EBT-PFX-088 | GGCTCACCTTCGGGTGGGCCTTTCTGCGTTACCGCATATGCTGGATCCTT | |
| EBT-PFX-089 | ACGTTGTAAAACGACGGCCAGTGCCGAGCTCTGCAGGTCGACTCTAGAG AAT | |
| EBT-PFX-090 | TTACCTTTTCATTCCCTACAGTTTTAGAGCTAGAAATAGC | gRNA01 |
| EBT-PFX-091 | TGTAGGGAATGAAAAGGTAAGCTAGCATTATACCTAGGAC | |
| EBT-PFX-133 | AAAAATTTGGAGTAAGGCAAGTTTTAGAGCTAGAAATAGC | gRNA02 |
| EBT-PFX-134 | TTGCCTTACTCCAAATTTTTGCTAGCATTATACCTAGGAC | |
| EBT-PFX-137 | GAAAAAATTTGGAGTAAGGCAAGTTTTAGAGCTAGAAATAGC | gRNA04 |
| EBT-PFX-138 | TTGCCTTACTCCAAATTTTTTCGCTAGCATTATACCTAGGAC | |
| EBT-PFX-139 | CAAATAGTAAAGACAGCTCCGTTTTAGAGCTAGAAATAGC | gRNA05 |
| EBT-PFX-140 | GGAGCTGTCTTTACTATTTGGCTAGCATTATACCTAGGAC | |
| EBT-PFX-141 | AATTGATCACTATCTGGGCAGTTTTAGAGCTAGAAATAGC | gRNA06 |
| EBT-PFX-142 | TGCCCAGATAGTGATCAATTGCTAGCATTATACCTAGGAC | |
| EBT-PFX-153 | ATCCATCCTATAATTCCTTCGTTTTAGAGCTAGAAATAGC | gRNA10 |
| EBT-PFX-154 | GAAGGAATTATAGGATGGATGCTAGCATTATACCTAGGAC | |
| EBT-PFX-155 | AACAAGTGGATGAAATTTTCGTTTTAGAGCTAGAAATAGC | gRNA11 |
| EBT-PFX-156 | GAAAATTTCATCCACTTGTTGCTAGCATTATACCTAGGAC | |
| EBT-PFX-178 | GGTCAGGGAATGCCAACTTAGTTTTAGAGCTAGAAATAGC | gRNA13 |
| EBT-PFX-179 | TAAGTTGGCATTCCCTGACCGCTAGCATTATACCTAGGAC | |
| EBT-PFX-180 | GATCAAGCAGATAAGATCAGGTTTTAGAGCTAGAAATAGC | gRNA14 |
| EBT-PFX-181 | CTGATCTTATCTGCTTGATCGCTAGCATTATACCTAGGAC | |
| EBT-PFX-182 | TTCAAATAGTAAAGACAGCTCCGTTTTAGAGCTAGAAATAGC | gRNA15 |
| EBT-PFX-183 | GGAGCTGTCTTTACTATTTGAAGCTAGCATTATACCTAGGAC | |
| EBT-PFX-201 | AATCAAGCAGATAAAATAAGGTTTTAGAGCTAGAAATAGC | gRNA19 |
| EBT-PFX-202 | CTTATTTTATCTGCTTGATTGCTAGCATTATACCTAGGAC | |

**Table S6** Primers used for the verification of base editing

| Primer | Sequence | Note |
|---|---|---|
| EBT-PFX-131 | AGGATAGGACATACCCTGTG | Verification of *pta* editing |
| EBT-PFX-132 | CATCTACAGACATGCCTGTTC | |
| EBT-PFX-130 | GGAGTAAGGCAAAGGAAGAC | Sequencing for *pta* |
| EBT-PFX-166 | TCCCCAATTTAGCATACTAGGC | Verification of *adhE1* editing |
| EBT-PFX-167 | CACATATGCCTCCAGTGCAT | |
| EBT-PFX-168 | TTACTGACTGCTCTGAGGCA | Sequencing for *adhE1* |
| EBT-PFX-169 | ATGCACTGGAGGCATATGTG | Verification of *adhE2* editing |
| EBT-PFX-170 | GTGCAACTCCAAGACTACCAT | |
| EBT-PFX-171 | AGGAGCACCAGCTTTAACTG | Sequencing for *adhE2* |
| EBT-PFX-163 | TGAAGAAGCGCTTCAAGTTC | Verification of *aor1* editing |
| EBT-PFX-164 | CTGCCTCTAATAGTGAATCTGC | |
| EBT-PFX-200 | CGTTGGTGCAGTTATGGGAT | Sequencing for *aor1* |
| EBT-PFX-187 | CTAAGGCAATGGGGATTGGA | Verification of *aor2* editing |
| EBT-PFX-188 | AGTTTCCACCTCCTTAGGCTA | |
| EBT-PFX-189 | GGGAGCAGAATTCAAAGCAG | Sequencing for *aor2* |

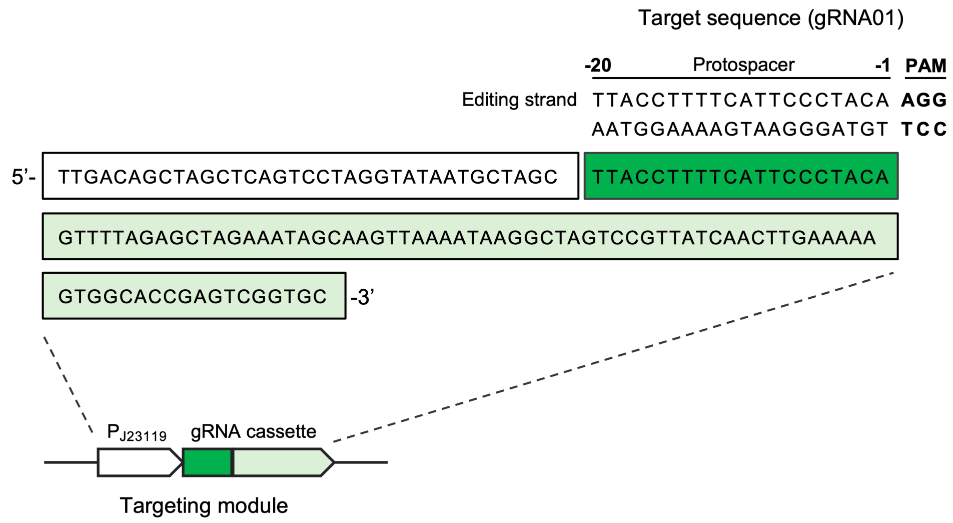


**Fig. S1** Scheme and sequence of gRNA cassette. Taking gRNA01 as an example, the sequences of $P_{J23119}$ promoter (white), gRNA01 (green), and gRNA scaffold of dCas9 from *S. pyogenes* (light green) are shown. For each line, the sequence reads from left to right and does not indicate the structure. Both strands of the target sequence are displayed. The gRNA01 will bind the complimentary strand of the editing strand, and potentially edits the Cs on the editing strand within the editing window. The nucleotide directly adjacent to the protospacer adjacent motifs (PAM) is counted as position -1 and the starting position.

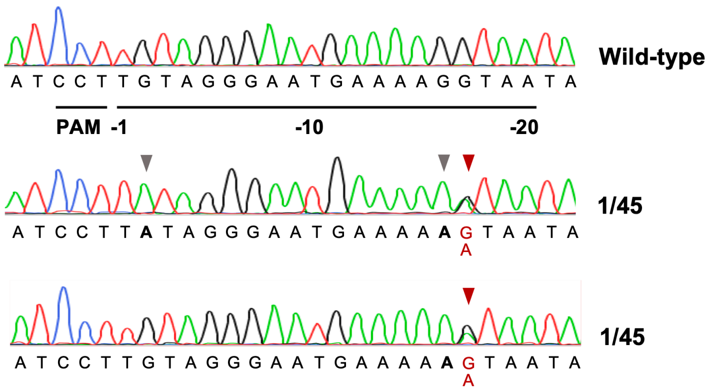


**Fig. S2** Mixed signals in the two colonies when editing *pta* using gRNA01**.** Grey arrows indicate clean mutations, and red arrows indicate mixed signals of G and A. (PAM: protospacer adjacent motifs.)

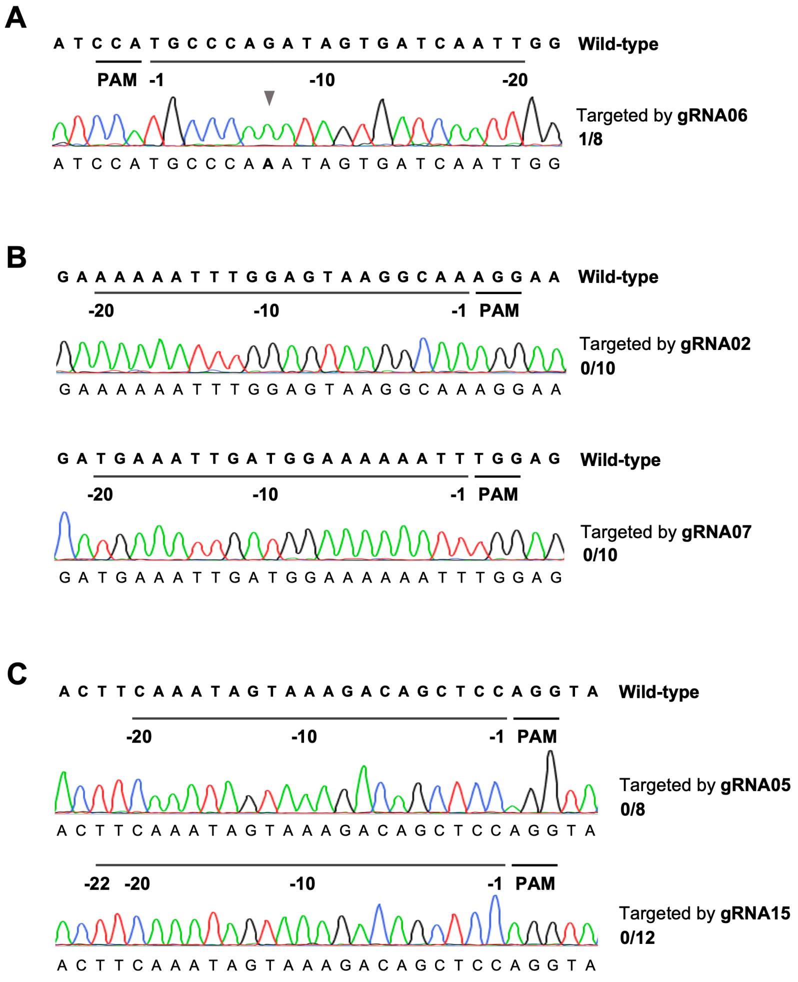


**Fig. S3** Editing principles. (A) Base editing in *pta* with gRNA06. Grey arrow indicates successful editing. (B) Evaluation of base editing on target sequences without Cs using gRNA02 and gRNA07. (C) Base editing with different length of gRNAs (gRNA05 and gRNA15) for editing the C at position -20. (PAM: protospacer adjacent motifs.)

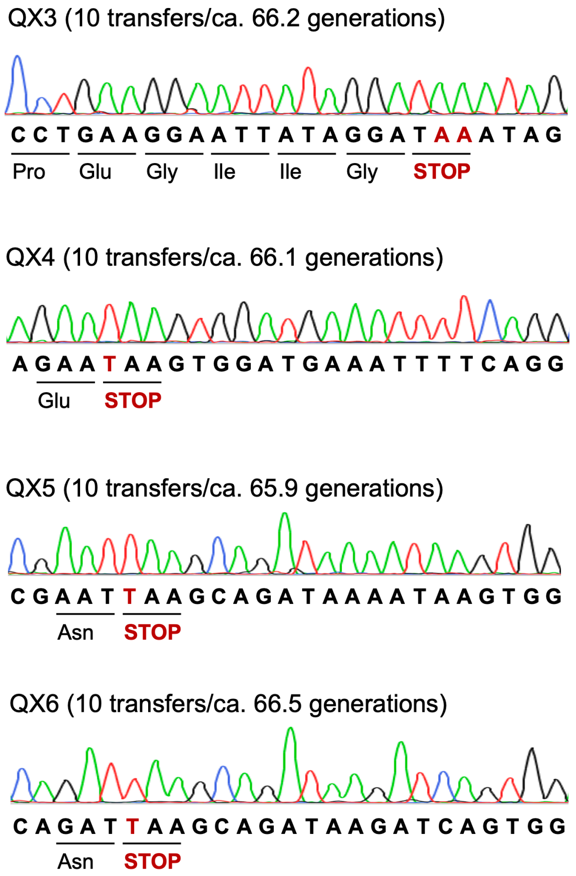


**Fig. S4** Stability test of four edited strains. QX3, QX4, QX5, and QX6 were transferred 10 times in RCM and the single-nucleotide variations were sequenced to check stability. The sequencing results of the 10th transfer are shown here.

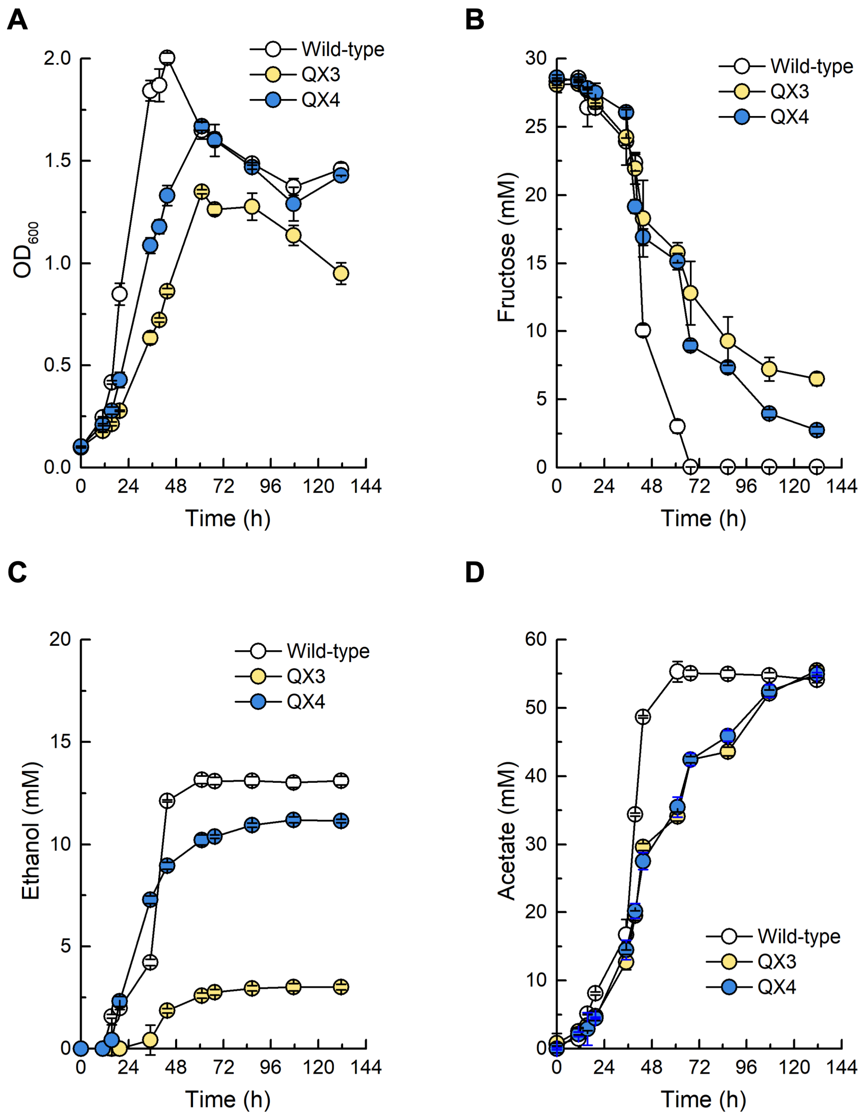


**Fig. S5** Fermentation performances of wild-type, QX3 (*adhE1* Trp169*), and QX4 (*adhE2* Gln33*) under heterotrophic conditions with 5 g/L (27.8 mM) of fructose as the carbon source. (A) Growth, (B) concentration of fructose, (C) concentrations of ethanol, and (D) concentration of acetate in the culture during fermentation.

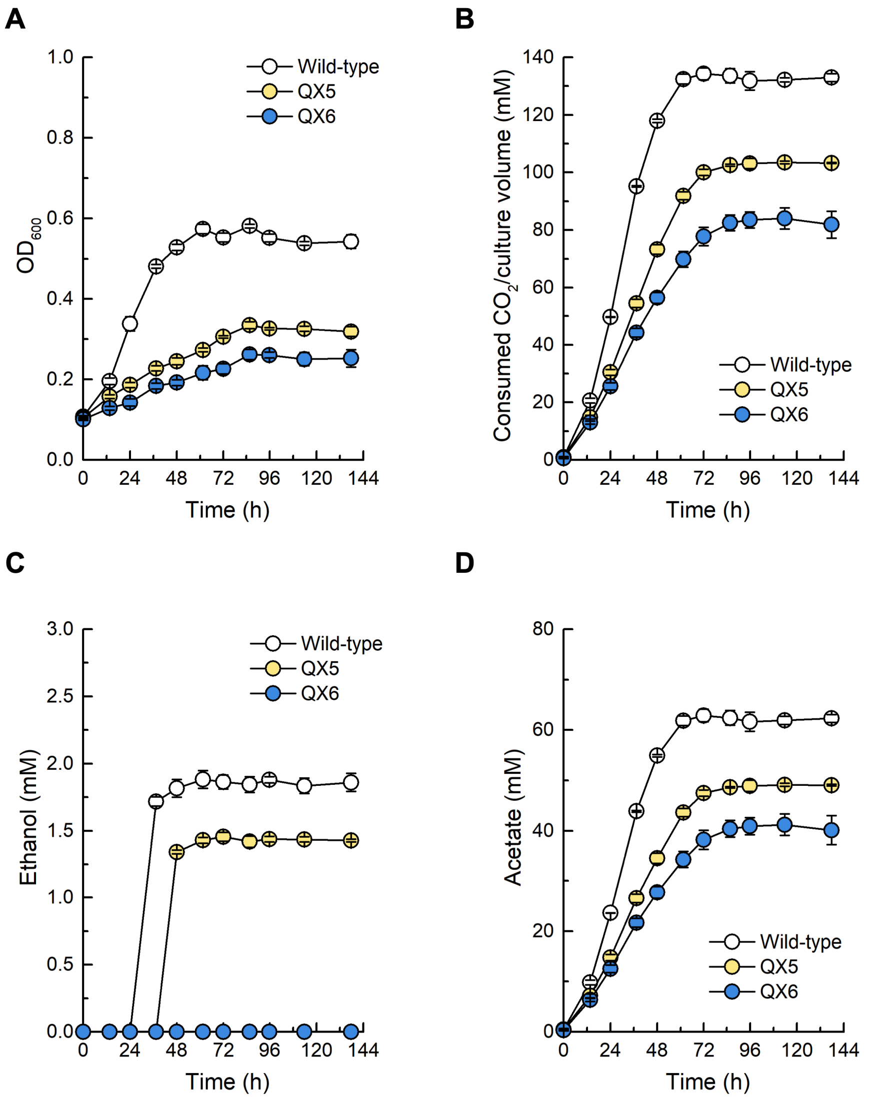


**Fig. S6** Fermentation performance of wild-type, QX5 (*aor1* Gln267*), and QX6 (*aor2* Gln267*) under autotrophic conditions with a gas mixture ($H_2$/$CO_2$, 80/20 vol-%, 1.5 bar) as the substrate. (A) Growth, (B) consumed $CO_2$ per culture volume, (C) concentrations of ethanol, and (D) concentration of acetate in the culture during fermentation.

### References


1. B. Molitor, K. Kirchner, A. W. Henrich, S. Schmitz, M. A. Rosenbaum, Expanding the molecular toolkit for the homoacetogen *Clostridium ljungdahlii*. *Sci. Rep.* **6**, 31518 (2016).
2. B. M. Woolston, D. F. Emerson, D. H. Currie, G. Stephanopoulos, Rediverting carbon flux in *Clostridium ljungdahlii* using CRISPR interference (CRISPRi). *Metab. Eng.* **48**, 243-253 (2018).
3. Y. Jiang *et al.*, Multigene editing in the *Escherichia coli* genome via the CRISPR-Cas9 system. *Appl. Environ. Microbiol.* **81**, 2506-2514 (2015).
4. S. Banno, K. Nishida, T. Arazoe, H. Mitsunobu, A. Kondo, Deaminase-mediated multiplex genome editing in *Escherichia coli*. *Nat. Microbiol.* **3**, 423-429 (2018).
5. C.-M. Klask, N. Kliem-Kuster, B. Molitor, L. T. Angenent, An open-source multiple-bioreactor system for replicable gas-fermentation experiments: Nitrate feed results in stochastic inhibition events, but improves ethanol production of *Clostridium ljungdahlii* with $CO_2$ and $H_2$. *bioRxiv* doi: 10.1101/2019.12.15.877050 (2019).
